# Making up your mind: Enhanced perceptual decision-making induced by stochastic resonance during non-invasive brain stimulation: Stochastic resonance in perceptual decision-making

**DOI:** 10.1101/175455

**Authors:** Onno van der Groen, Matthew F. Tang, Nicole Wenderoth, Jason B. Mattingley

## Abstract

Perceptual decision-making relies on the gradual accumulation of noisy sensory evidence until a specified boundary is reached and an appropriate response is made. It might be assumed that adding noise to a stimulus, or to the neural systems involved in its processing, would interfere with the decision process. But it has been suggested that adding an optimal amount of noise can, under appropriate conditions, enhance the quality of subthreshold signals in nonlinear systems, a phenomenon known as *stochastic resonance*. Here we asked whether perceptual decisions obey these stochastic resonance principles by adding noise directly to the visual cortex using transcranial random noise stimulation (tRNS) while participants judged the direction of motion in foveally presented random-dot motion arrays. Consistent with the stochastic resonance account, we found that adding tRNS bilaterally to visual cortex enhanced decision-making when stimuli were just below, but not well below or above, perceptual threshold. We modelled the data under a drift diffusion framework to isolate the specific components of the multi-stage decision process that were influenced by the addition of neural noise. This modelling showed that tRNS increased drift rate, which indexes the rate of evidence accumulation, but had no effect on bound separation or non-decision time. These results were specific to bilateral stimulation of visual cortex; control experiments involving unilateral stimulation of left and right visual areas showed no influence of random noise stimulation. Our study is the first to provide causal evidence that perceptual decision-making is susceptible to a stochastic resonance effect induced by tRNS, and that this effect arises from selective enhancement of the rate of evidence accumulation for sub-threshold sensory events.

## Results and Discussion

Noise is an intrinsic property of all biological systems [2]. Typically, noise is viewed as being detrimental for neuronal computations and the behaviors they regulate [2, 3], including decision-making [4]. A key limiting factor in decision-making arises from noisy representations of sensory evidence in the brain [5, 6]. On this view, noisy sensory information representations are not optimal, and this leads to errors in decisions. However, small amounts of noise added to a nonlinear system can increase the stimulus quality by increasing the signal-to-noise ratio (SNR)[7]. This phenomenon is known as *stochastic resonance*, and its expression has been demonstrated in different sensory modalities [8-10]. Stochastic resonance occurs when an optimal amount of noise is added to a sub-threshold signal, which makes the signal cross a decision threshold, and therefore enhances detection performance (Figure 1).

**Figure 1:**
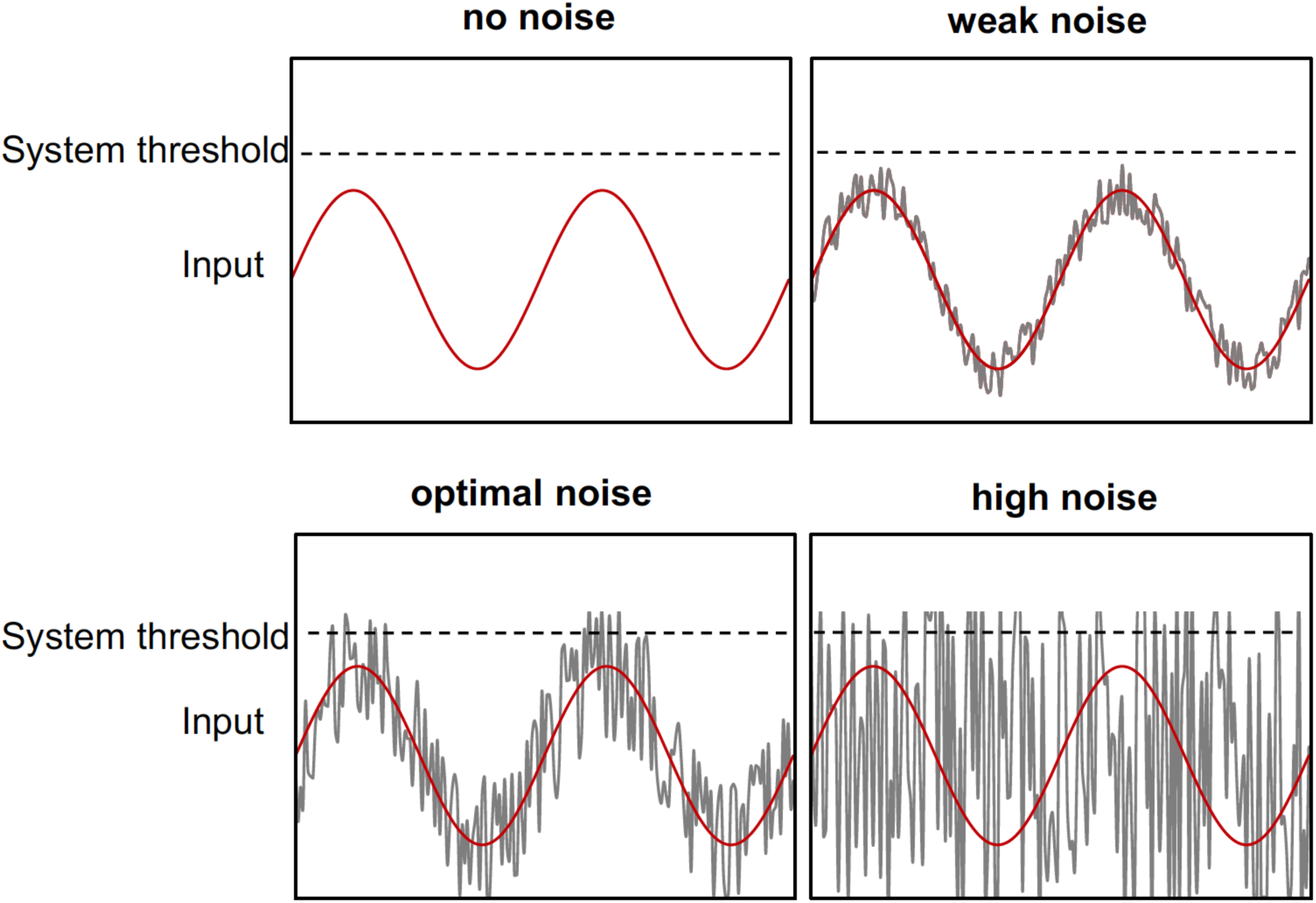
Stochastic resonance occurs when an optimal level of noise is added to a subthreshold signal. In this example the signal alone (red sinusoid) remains below the perceptual threshold (dotted line). Adding an optimal amount of noise (grey line) periodically raises the stimulus above the system threshold. If the added noise is too weak, the threshold is not crossed. Conversely, if the noise is too strong the signal remains buried and cannot be discriminated from the noise[1].

In a typical stochastic resonance experiment, predefined noise intensities are added to a signal. Adding too little noise does not cause threshold crossings when the signal is well below the detection threshold, and performance remains unaffected. By contrast, when too much noise is added the signal gets buried in the noise and performance declines [11-14]. This results in an ‘inverted U’ relationship between noise intensity and detection performance, which is a key signature of the stochastic resonance effect [8-10]. At the neurophysiological level, it has been shown that adding an optimal amount of noise to a subthreshold signal pushes otherwise silent sensory neurons above the spiking threshold [10, 15, 16]. A common way of adding noise in a stochastic resonance context is to add it directly to the sensory stimulus. In that case, the noise could increase peripheral receptor sensitivity [17], which does not permit investigation of whether central neural processes in decision-making are sensitive to a stochastic resonance mechanism. Recently, we showed it is possible to induce a stochastic resonance effect in a simple detection task when noise is added to the cortex directly with tRNS [18]. In that study, participants had to detect a weak visual stimulus which was either sub-or suprathreshold. On each trial, tRNS with a predefined intensity was added to the visual cortex. We demonstrated that adding an optimal noise level with tRNS significantly enhanced detection performance for subthreshold, but not suprathreshold, stimuli.

Here we asked whether perceptual decision-making, as opposed to mere detection, can be enhanced for subthreshold visual stimuli by the application of tRNS over visual cortex, either bilaterally (Experiment 1) or unilaterally over the left and right visual cortex (Experiments 2 and 3). To examine these effects, we used drift diffusion modeling (DDM, see Figure 2A) [19, 20] to uncover what aspects of the decision process were influenced by tRNS. Under the DDM framework, decision-making requires the collection of noisy sensory evidence over time. The collection of sensory evidence continues until a criterion is reached, and based on the collected sensory evidence, a decision is made [21, 22]. The DDM is a highly successful approach that has been used over a wide range of tasks [23].

**Figure 2:**
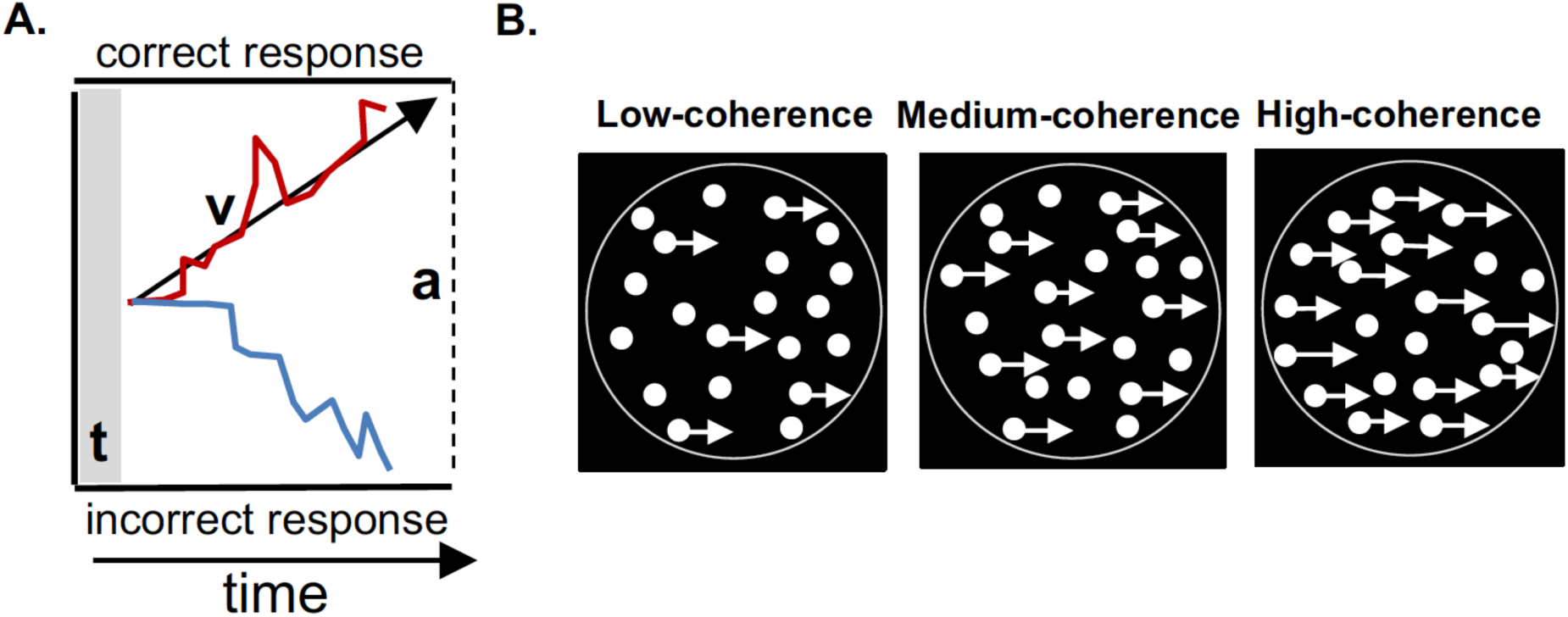
**A:** Schematic of the Drift Diffusion Modelling (DDM) framework used to model perceptual decision-making in the dot motion task. In the model, evidence is accumulated over time until a response boundary is crossed. *t* is the non-decision time, which includes the time taken to execute a motor response. *v* is the drift rate, which reflects the rate at which sensory evidence is accumulated. This parameter is taken as an index of the quality of sensory information. *a* represents the boundary separation (*correct* at the top, *incorrect* at the bottom), indicating how much information is needed to make a decision. **B:** Schematic of the random dot-motion task in which participants judged whether signal dots moved on average to the left or right. Task difficulty was titrated by altering the proportion of coherently moving dots (shown with arrows attached, for purposes of illustration) amongst randomly moving dots. In this example the coherent motion is rightward, but in the experiment the dots were equally likely to move toward the left or right. For display purposes, we depicted here the outline of the imaginary circle in which the dots were presented.

We used a random-dot-motion task (RDM-task) as the perceptual decision-making task of choice (see Supplemental Information). The RDM-task is widely used in studies of perceptual decision-making, and has well characterized neural correlates [24, 25]. Participants fixated on a centrally presented array of randomly moving dots within which a proportion of the dots moved coherently in a common direction (leftward or rightward; see Figure 2B). Participants judged the common direction of movement (two-alternative forced-choice/2-AFC) as quickly and accurately as possible. The difficulty of the task was parametrically manipulated by altering the proportion of signal dots that moved coherently in a given trial (3%, 6%, 12%, 25% or 50% coherence). The benefit of this task is that it allows for the continuous accumulation of sensory evidence over a period of several hundred milliseconds, which facilitates investigation of the underlying processes involved in decision-making [21, 26, 27].

If the stochastic resonance model applies to perceptual decision-making, then the addition of relatively small amounts of noise should enhance motion discrimination performance. The added noise will likely increase the quality of the sensory evidence for coherent motion trials in which the signal is just below threshold, but not for trials in which the signal is well below or above threshold. Conversely, the addition of large amounts of noise should either have no effect on perceptual thresholds, or should impair performance slightly for displays at or above threshold [8]. We therefore applied four different tRNS intensities (0.25, 0.375, 0.5 and 0.75 mA; 100-640 Hz zero-mean Gaussian white noise) while participants engaged in the RDM-task. These tRNS intensities result in current densities that we have shown previously are able to induce a stochastic resonance effect in a visual contrast detection task [18]. The tRNS was applied in blocks of 20 trials at one of the four intensities, with each block of stimulation followed by a 20-trial block of no-stimulation to minimise build-up of any cumulative effects of the stimulation. Participants were blinded to the tRNS conditions. Consistent with several previous investigations, no participant reported awareness of the stimulation during de-briefing [28, 29].

## Experiment 1: Effect of bilateral visual cortex stimulation on perceptual decision-making

In Experiment 1, we stimulated visual cortex bilaterally with tRNS in 15 participants (see Figure 3 and Figure S2). The coherence levels of 3% and 6% were subthreshold (average detection performance < 0.63%), i.e., performance was below the detection threshold, which corresponded to 75% correct in our task. For the analysis, we calculated the group %correct-choice-index (%CCI) for each coherence level and each tRNS intensity by dividing the %correct motion-direction responses under tRNS by the %correct responses when no tRNS was applied (baseline), as given in the following formula:

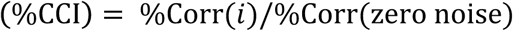

where *i* denotes each of the 4 tested noise intensities. As shown in the left panel in Figure 3, for the 6% coherence condition, which was just below threshold in the no-tRNS condition, motion discrimination performance improved when tRNS was applied at a relatively low intensity, whereas performance remained unaffected for the other coherence levels and noise intensities. To quantify these effects, we performed a 4 (tRNS intensity) x 5 (coherence level) within-subjects ANOVA on the %CCI data. There was a significant interaction between coherence level and tRNS-intensity (F(12,156) = 2.47 p < 0.01, Cohen’s f= 0.43). To isolate the source of this interaction, one-way ANOVAs were conducted for each coherence level separately. For the 6% coherence condition only (red symbols in Figure 3), performance was significantly affected by the different tRNS intensities (F(3,39) = 3.56 p = 0.02 Cohen’s f=0.52). There were no other significant main effects or interactions for the coherence conditions of 3%, 12%, 25% or 50%. Post-hoc tests were conducted to compare performance in the 6% coherence condition at each noise level against the baseline. All p-values were corrected for multiple comparisons. These comparisons revealed that a tRNS intensity of 0.25mA significantly enhanced motion discrimination performance relative to baseline (t(13) = 3.39 p_corrected_ < 0.02). A similar enhancement was evident for the 6% coherence level at an intensity of .375mA, but this effect did not survive our stringent correction for multiple comparisons, (t(13) = 2.53, p_corrected_ > 0.1). These results suggest that perceptual decision-making for sensory stimuli that are just below threshold can be improved by adding a small amount of neural noise over bilateral visual cortex, consistent with predictions arising from the stochastic resonance principle [8].

**Figure 3:**
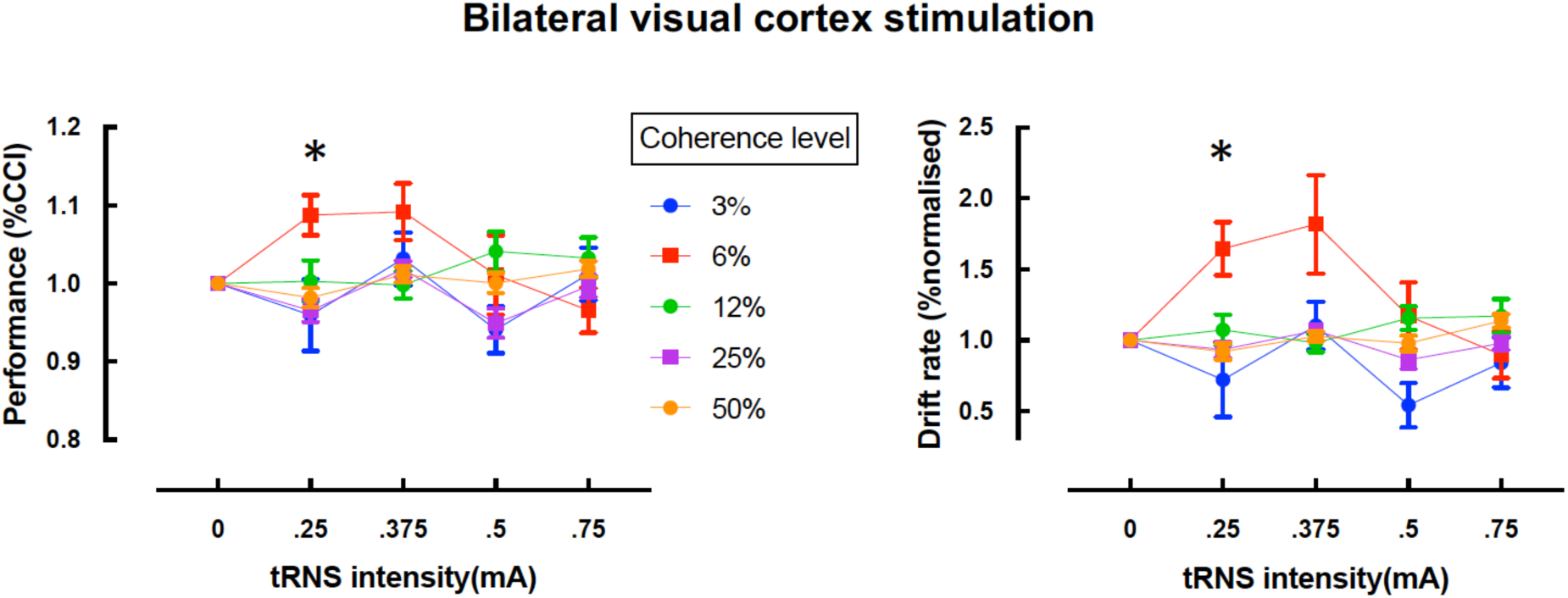
Effects of transcranial random noise stimulation (tRNS) on perceptual decision-making in the dot-motion discrimination task for bilateral stimulation. The left panel shows performance for each motion coherence level as a function of tRNS intensity. The right panel shows the drift rate derived from modelling of the data shown in the corresponding plot to the left. *p_corrected_ < 0.05.

Next we employed the drift diffusion framework to accurately model the processes involved in decision-making based on the accuracy and response time data obtained from the decision-making task. Specifically, we used the hierarchical drift diffusion model (HDDM, [30]) to determine which aspect of decision-making was affected by tRNS. We normalized the obtained DDM-parameters relative to the zero noise condition in the same way as the behavioral data, as described above. As shown in the right panel of Figure 3, the drift rate was markedly affected by tRNS for the 6% coherence condition, whereas it appears to be unaffected for the remaining coherence levels. We submitted the drift-rate parameter to a 5 x 4 repeated measures ANOVA. This analysis revealed a significant main effect of tRNS-intensity (F(3,39) = 2.85, p = 0.049) and of coherence level (F(4,52) = 3.18, p = 0.02 on drift rate, as well as a significant tRNS-intensity x coherence level interaction (F(12,156) = 3.22, p < .01, Cohen’s f = 0.47). To isolate the source of the significant interaction, one-way ANOVAs were conducted for each coherence level separately. Consistent with the behavioral data, there was a significant effect of tRNS intensity on drift rate in the 6% coherence condition (F(3,39) = 5.63, p < .01, Cohen’s f = .58), but no significant effects for the other coherence levels (3%, 12%, 25%, 50%).

Post-hoc tests were conducted to compare performance in the 6% coherence condition against the baseline for each noise level. For the tRNS intensity of .25mA, the drift rate for the 6% coherence condition was significantly higher than baseline (t(13) = 3.44, p_corrected_ < 0.02, corrected for multiple-comparisons). A similar benefit for the 6% coherence condition was apparent for the tRNS intensity of .375mA, but this effect did not survive correction for multiple comparisons (t(13) = 2.55, p = 0.1). Separate 5 x 4 repeated measures ANOVAs revealed no significant effects for the bound-separation parameter (all p > 0.06), and no significant effects for non-decision time (all p > 0.13).

Previous studies of visual motion discrimination have shown reliable effects of offline transcranial electrical stimulation – as opposed to the online effects reported here – following unilateral stimulation of left or right visual cortex in isolation [31-33]. We therefore conducted two further experiments to determine whether the stochastic resonance effects we observed for bilateral tRNS in Experiment 1 also arise for unilateral visual stimulation. We also modelled the current spread for the electrode montage used in each experiment using the SimNibs toolbox [34] (Figure 4).

**Figure 4:**
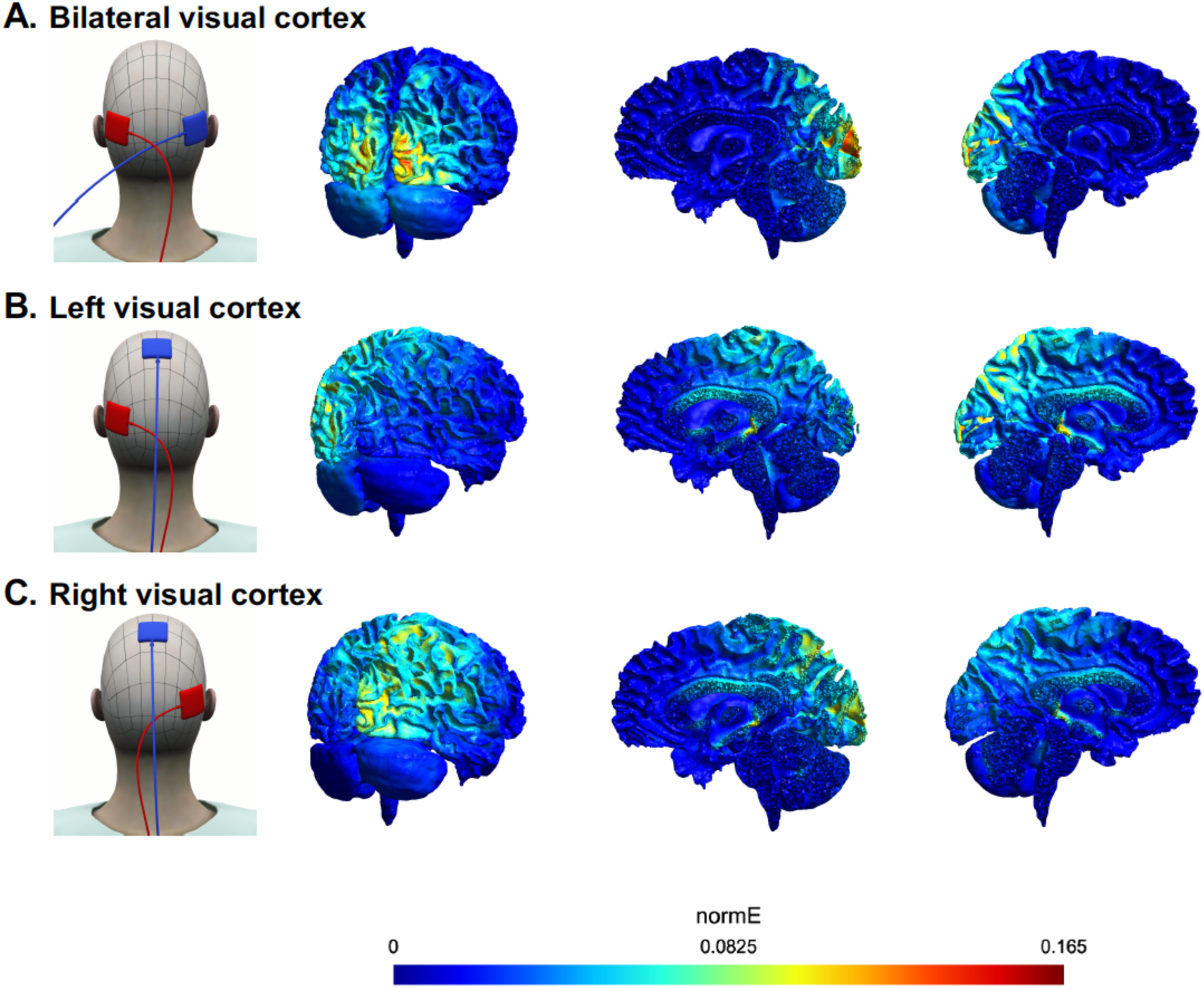
Electrode pad montages and modelled electrical field strength (normE) for each of the three tRNS experiments. **A.** Bilateral visual cortex stimulation (Experiment 1). **B.** Left visual cortex stimulation (Experiment 2). **C.** Right visual cortex stimulation (Experiment 3).

## Experiments 2 and 3 – Effect of unilateral visual cortex stimulation on perceptual decision-making

In Experiments 2 and 3 we applied tRNS unilaterally to the left and right visual cortex, respectively, using two new groups of 15 participants each. Figures 5A and 5B show the behavioral results for these two experiments. Neither left nor right unilateral tRNS produced the characteristic inverted-U tuning curve observed in Experiment 1 for the 6% coherence condition during bilateral stimulation. To characterize the data statistically, we employed the same analytic approach as for the bilateral tRNS experiment, for both the behavioural data and the drift diffusion modelling. There was no significant interaction between stimulation intensity and coherence level for either left unilateral or right unilateral visual cortex stimulation (p > .05 for all key comparisons). Thus, there was no evidence for the stochastic resonance effect observed during bilateral stimulation in experiment 1.

**Figure 5:**
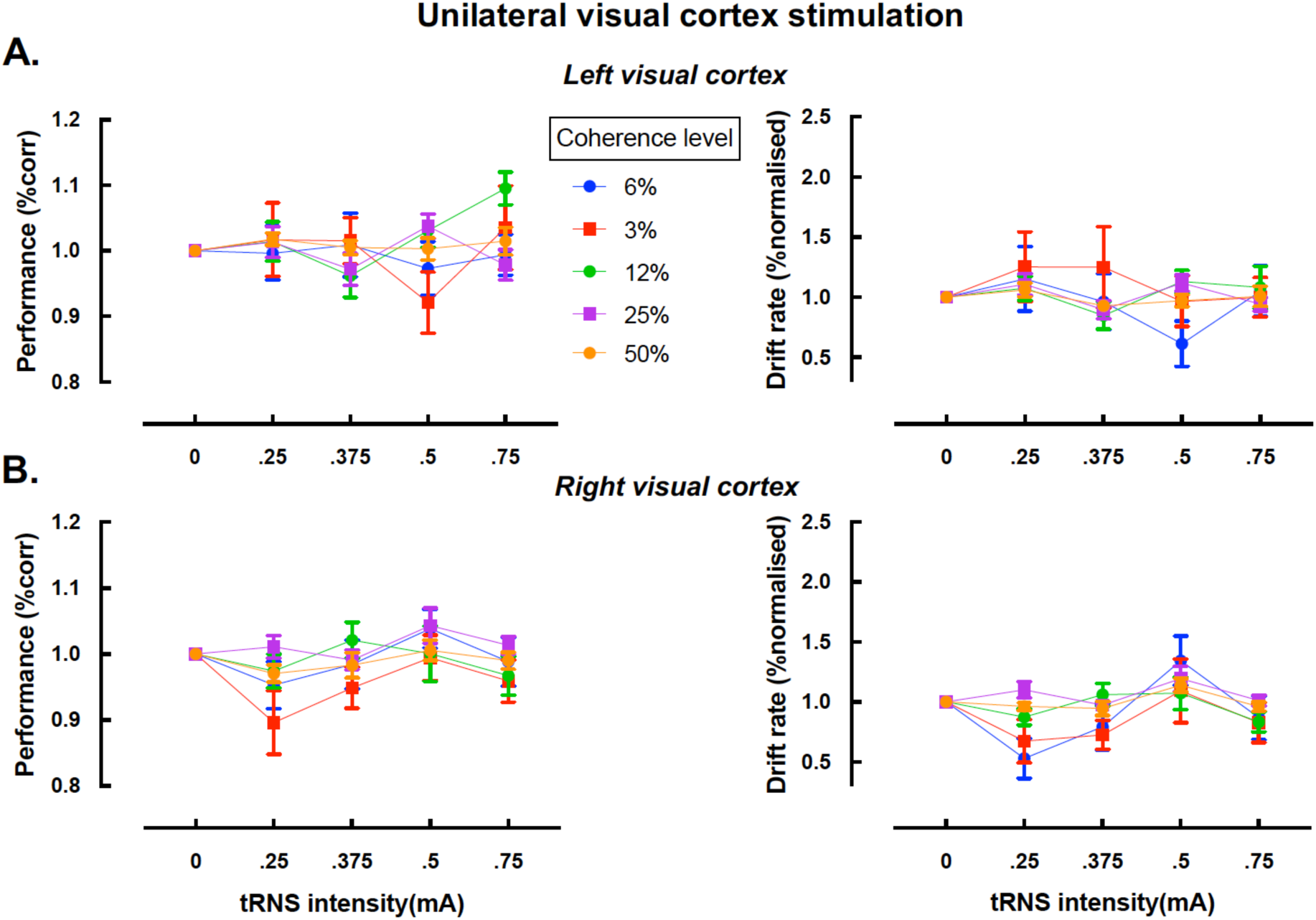
Effects of transcranial random noise stimulation (tRNS) on perceptual decision-making in the dot-motion discrimination task for unilateral stimulation for the left visual cortex **(A)** and right visual cortex **(B)** stimulation. The left panels show performance for each motion coherence level as a function of tRNS intensity. The right panels show the drift rate derived from modelling of the data shown in the corresponding plots to the left.

Analysis of the baseline data in all three experiments revealed no interaction between coherence level and tRNS intensity (repeated measures ANOVA with within-subjects factor of coherence level and between-subjects factor of experiment, F(2,39) = 1.15, p > .32), suggesting that the stochastic resonance-effect observed in Experiment 1 was not driven by differences in baseline performance between the three experiments. Across all three experiments, there was a highly significant main effect of coherence level on performance, as expected. For completeness, we also report here a small number of significant main effects which are not related to the central stochastic resonance hypothesis under examination in this study (see also Figure S2). First, there was a small but consistent main effect of tRNS intensity on accuracy during right visual cortex stimulation, F(3,39) = 3.13 p = .036, Cohen’s f =0.49. Post-hoc contrasts revealed that this effect was driven by overall poorer performance for the .25mA tRNS intensity, regardless of motion coherence level, t(69) = −2.78 p_corrected_ < 0.03. This decrease in performance was mirrored by a significant main effect of tRNS-intensity on drift rate (see Figure 5B and Table S1), (F(3,39) = 4.54 p < .01, Cohen’s f = 0.59, which was again specific to the .25mA tRNS intensity, (t(69) = 2.67 p_corrected_ = .04), regardless of motion coherence level. Second, there was a significant main effect of coherence level on bound-separation during stimulation of the right visual cortex, F(4,52) = 3.09 p = .024, Cohen’s f = 0.4 (see Supplemental Information). Post-hoc tests showed that the bounds were significantly closer together for the highest (50%) coherence condition, t(55) = 3.16 p_corrected_ < .04, relative to baseline), but there were no significant effects on bound separation for the other coherence levels.

## Conclusions

We have shown that adding a small amount of noise bilaterally to the visual cortex can enhance perceptual decision-making in a motion discrimination task, particularly for subthreshold stimuli (6% coherence). When modeled as a drift-diffusion process, this tRNS-induced performance improvement coincided with an increase in the rate of evidence accumulation, reflected as a change in the model’s drift-rate parameter. The same model revealed no change in either bound-separation or non-decision time, suggesting that an optimal level of neural noise exclusively improves perceptual decision-making by enhancing sensory information quality, consistent with a stochastic resonance mechanism ([8-10], see Supplemental Information for the model fits). In line with previous work [18], we showed that the stochastic resonance effect was strongest when appropriate tRNS intensities were added to the 6% coherence condition, i.e. to a *subthreshold* stimulus, as indicated by the average baseline detection accurarcy of 60%. Note that all tRNS intensities and coherence levels were randomized over participants to account for any aftereffects, fatigue or learning effects across conditions.

There was no evidence for a stochastic resonance effect when noise was applied unilaterally to the visual cortex. This absence of a performance-enhancing effect for unilateral tRNS was not due to differences in baseline performance between the groups: detection performance in the 6% coherence condition was similar across experiments (Experiment 1 – 60%; Experiment 2 – 58%; Experiment 3 – 57%). Modelling of the electrical field for each electrode montage (Figure 4) indicated a higher peak current when the tRNS was applied bilaterally than in the unilateral stimulation conditions. It is unlikely that this apparent difference in current densities prevented a stochastic resonance effect for the unilateral stimulation conditions, however, because the same absolute current densities during bilateral stimulation were also reached during unilateral stimulation but at higher tRNS intensities.

The visual stimuli employed in our motion discrimination task were always presented in the centre of the display, and thus would have been processed initially by visual areas in both the left and right hemsipheres [35]. Given that unilateral visual cortex stimulation did not influence motion-discrimination performance, it is most parsimonious to conclude that visual areas in *both* hemispheres must be stimulated concurrently with tRNS for the stochastic resonance effect to occur. Because of the relatively diffuse nature of transcranial electrical stimulation in general [36], it is not possible to determine the specific anatomical regions that mediate the stochastic resonance effect we observed. The primary visual cortex (V1) [37] and motion area V5/MT are both crucial for the processing of dynamically moving visual stimuli [38-40]. These two areas are highly interconnected, so our bilateral stimulation protocol might have impacted motion processing in area V5/MT, enhanced signal quality in area V1, or both. Further work using more focal stimulation techniques (e.g., transcranial magnetic stimulation) will be needed to pinpoint the visual areas responsible for the stochastic resonance effects we report here.

Our results are in line with recent work that employed a similar task to show that decision-making is sensitive to the addition of noise to visual motion stimuli [41]. Critically, our findings extend these results by demonstrating that a stochastic resonance effect can be induced in a decision-making task when noise is directly applied to the visual cortex with tRNS [42, 43]. Moreover, we are the first to show that this stochastic resonance effect enhances the quality of information processing as indicated by an accelerated rate of evidence accumulation. The underlying mechanism for the observed tRNS effect is not completely understood. However, single unit recordings have shown that sensory neurons in the visual cortex are sensitive to a stochastic resonance mechanism, e.g., there is an increase in the SNR of the firing rate of neurons when an optimal level of noise is applied to a visual stimulus [44]. This is likely due to the recruitment of voltage-gated sodium channels by the noise [45-47].

## References

1. Davis, G., and Plaisted-Grant, K. (2015). Low endogenous neural noise in autism. Autism : the international journal of research and practice 19, 351–362.

2. Tsimring, L.S. (2014). Noise in biology. Rep Prog Phys 77, 026601.

3. Faisal, A.A., Selen, L.P., and Wolpert, D.M. (2008). Noise in the nervous system. Nature reviews. Neuroscience 9, 292–303.

4. Heekeren, H.R., Marrett, S., Bandettini, P.A., and Ungerleider, L.G. (2004). A general mechanism for perceptual decision-making in the human brain. Nature 431, 859–862.

5. Kaufman, M.T., and Churchland, A.K. (2013). Cognitive neuroscience: sensory noise drives bad decisions. Nature 496, 172–173.

6. Brunton, B.W., Botvinick, M.M., and Brody, C.D. (2013). Rats and Humans Can Optimally Accumulate Evidence for Decision-Making. Science 340, 95–98.

7. Gammaitoni, L., Hänggi, P., Jung, P., and Marchesoni, F. (1998). Stochastic resonance. Reviews of modern physics 70, 223.

8. Moss, F. (2004). Stochastic resonance and sensory information processing: a tutorial and review of application. Clinical Neurophysiology 115, 267–281.

9. McDonnell, M.D., and Abbott, D. (2009). What is stochastic resonance? Definitions, misconceptions, debates, and its relevance to biology. PLoS Comput Biol 5, e1000348.

10. Hanggi, P. (2002). Stochastic resonance in biology - How noise can enhance detection of weak signals and help improve biological information processing. Chemphyschem 3, 285–290.

11. Collins, J.J., Imhoff, T.T., and Grigg, P. (1996). Noise-enhanced tactile sensation. Nature 383, 770–770.

12. Simonotto, E., Riani, M., Twitty, J., and Moss, F. (1997). Human perception of subthreshold, noise-enhanced visual images. Fr Art Int 37, 176–183.

13. Zeng, F.G., Fu, Q.J., and Morse, R. (2000). Human hearing enhanced by noise. Brain Res 869, 251–255.

14. Lugo, E., Doti, R., and Faubert, J. (2008). Ubiquitous crossmodal Stochastic Resonance in humans: auditory noise facilitates tactile, visual and proprioceptive sensations. Plos One 3, e2860.

15. Douglass, J.K., Wilkens, L., Pantazelou, E., and Moss, F. (1993). Noise enhancement of information transfer in crayfish mechanoreceptors by stochastic resonance. Nature 365, 337–340.

16. Wiesenfeld, K., Pierson, D., Pantazelou, E., Dames, C., and Moss, F. (1994). Stochastic resonance on a circle. Phys Rev Lett 72, 2125.

17. Mendez-Balbuena, I., Manjarrez, E., Schulte-Monting, J., Huethe, F., Tapia, J.A., Hepp-Reymond, M.C., and Kristeva, R. (2012). Improved sensorimotor performance via stochastic resonance. The Journal of neuroscience : the official journal of the Society for Neuroscience 32, 12612–12618.

18. van der Groen, O., and Wenderoth, N. (2016). Transcranial Random Noise Stimulation of Visual Cortex: Stochastic Resonance Enhances Central Mechanisms of Perception. The Journal of neuroscience : the official journal of the Society for Neuroscience 36, 5289–5298.

19. Ratcliff, R. (1978). A theory of memory retrieval. Psychological review 85, 59.

20. Ratcliff, R., and McKoon, G. (2008). The diffusion decision model: theory and data for two-choice decision tasks. Neural Comput 20, 873–922.

21. Gold, J.I., and Shadlen, M.N. (2007). The neural basis of decision making. Annu Rev Neurosci 30, 535–574.

22. Braun, J., and Mattia, M. (2010). Attractors and noise: twin drivers of decisions and multistability. Neuroimage 52, 740–751.

23. Forstmann, B.U., Ratcliff, R., and Wagenmakers, E.J. (2016). Sequential Sampling Models in Cognitive Neuroscience: Advantages, Applications, and Extensions. Annu Rev Psychol 67, 641–666.

24. Shadlen, M.N., and Newsome, W.T. (2001). Neural basis of a perceptual decision in the parietal cortex (area LIP) of the rhesus monkey. Journal of neurophysiology 86, 1916–1936.

25. Ferrera, V.P., Rudolph, K.K., and Maunsell, J.H. (1994). Responses of neurons in the parietal and temporal visual pathways during a motion task. The Journal of neuroscience : the official journal of the Society for Neuroscience 14, 6171–6186.

26. Newsome, W.T., Britten, K.H., and Movshon, J.A. (1989). Neuronal correlates of a perceptual decision. Nature 341, 52–54.

27. Britten, K.H., Shadlen, M.N., Newsome, W.T., and Movshon, J.A. (1992). The analysis of visual motion: a comparison of neuronal and psychophysical performance. The Journal of neuroscience : the official journal of the Society for Neuroscience 12, 4745–4765.

28. Ambrus, G.G., Paulus, W., and Antal, A. (2010). Cutaneous perception thresholds of electrical stimulation methods: Comparison of tDCS and tRNS. Clinical Neurophysiology 121, 1908–1914.

29. Fertonani, A., Pirulli, C., and Miniussi, C. (2011). Random noise stimulation improves neuroplasticity in perceptual learning. The Journal of neuroscience : the official journal of the Society for Neuroscience 31, 15416–15423.

30. Wiecki, T.V., Sofer, I., and Frank, M.J. (2013). HDDM: Hierarchical Bayesian estimation of the Drift-Diffusion Model in Python. Front Neuroinform 7, 14.

31. Zito, G.A., Senti, T., Cazzoli, D., Muri, R.M., Mosimann, U.P., Nyffeler, T., and Nef, T. (2015). Cathodal HD-tDCS on the right V5 improves motion perception in humans. Front Behav Neurosci 9, 257.

32. Antal, A., Varga, E.T., Nitsche, M.A., Chadaide, Z., Paulus, W., Kovacs, G., and Vidnyanszky, Z. (2004). Direct current stimulation over MT+/V5 modulates motion aftereffect in humans. Neuroreport 15, 2491–2494.

33. Kar, K., and Krekelberg, B. (2014). Transcranial alternating current stimulation attenuates visual motion adaptation. The Journal of neuroscience : the official journal of the Society for Neuroscience 34, 7334–7340.

34. Thielscher, A., Antunes, A., and Saturnino, G.B. (2015). Field modeling for transcranial magnetic stimulation: A useful tool to understand the physiological effects of TMS? Conf Proc IEEE Eng Med Biol Soc 2015, 222–225.

35. d'Avossa, G., Tosetti, M., Crespi, S., Biagi, L., Burr, D.C., and Morrone, M.C. (2007). Spatiotopic selectivity of BOLD responses to visual motion in human area MT. Nature neuroscience 10, 249–255.

36. Woods, A.J., Antal, A., Bikson, M., Boggio, P.S., Brunoni, A.R., Celnik, P., Cohen, L.G., Fregni, F., Herrmann, C.S., Kappenman, E.S., et al. (2016). A technical guide to tDCS, and related non-invasive brain stimulation tools. Clinical neurophysiology : official journal of the International Federation of Clinical Neurophysiology 127, 1031–1048.

37. Ajina, S., Kennard, C., Rees, G., and Bridge, H. (2015). Motion area V5/MT+ response to global motion in the absence of V1 resembles early visual cortex. Brain 138, 164–178.

38. Salzman, C.D., Murasugi, C.M., Britten, K.H., and Newsome, W.T. (1992). Microstimulation in visual area MT: effects on direction discrimination performance. The Journal of neuroscience : the official journal of the Society for Neuroscience 12, 2331–2355.

39. Thakral, P.P., and Slotnick, S.D. (2011). Disruption of MT impairs motion processing. Neuroscience letters 490, 226–230.

40. Newsome, W.T., and Pare, E.B. (1988). A selective impairment of motion perception following lesions of the middle temporal visual area (MT). The Journal of neuroscience : the official journal of the Society for Neuroscience 8, 2201–2211.

41. Trevino, M., De la Torre-Valdovinos, B., and Manjarrez, E. (2016). Noise Improves Visual Motion Discrimination via a Stochastic Resonance-Like Phenomenon. Frontiers in human neuroscience 10, 572.

42. Miniussi, C., Harris, J.A., and Ruzzoli, M. (2013). Modelling non-invasive brain stimulation in cognitive neuroscience. Neuroscience and biobehavioral reviews 37, 1702–1712.

43. Schwarzkopf, D.S., Silvanto, J., and Rees, G. (2011). Stochastic resonance effects reveal the neural mechanisms of transcranial magnetic stimulation. The Journal of neuroscience : the official journal of the Society for Neuroscience 31, 3143–3147.

44. Funke, K., Kerscher, N.J., and Worgotter, F. (2007). Noise-improved signal detection in cat primary visual cortex via a well-balanced stochastic resonance-like procedure. European Journal of Neuroscience 26, 1322–1332.

45. Onorato, I., D'Alessandro, G., Di Castro, M.A., Renzi, M., Dobrowolny, G., Musaro, A., Salvetti, M., Limatola, C., Crisanti, A., and Grassi, F. (2016). Noise Enhances Action Potential Generation in Mouse Sensory Neurons via Stochastic Resonance. Plos One 11, e0160950.

46. Schoen, I., and Fromherz, P. (2008). Extracellular stimulation of mammalian neurons through repetitive activation of Na+ channels by weak capacitive currents on a silicon chip. Journal of neurophysiology 100, 346–357.

47. Bromm, B. (1968). Die Natrium-Gleichrichtung der unterschwellig erregten Membran in der quantitativen Formulierung der Ionentheorie. Pflügers Archiv 302, 233–244.

